# ECCO: A Python UI for Performing Ensemble Clustering Combined with Cluster Optimization on Omics Data

**DOI:** 10.1101/2022.11.03.515009

**Authors:** Brady David Hislop, Kenna Brown, Hunter Hasskamp, Connor Boone, Mark Greenwood, Chelsea M. Heveran, Ronald K. June

## Abstract

Modern biological research often leverages clustering to elucidate disease endotypes and underlying mechanisms. Mono-cluster solutions remain the predominant method; however, this approach has concerning pitfalls, motivating the need for ensemble clustering methods. We present Ensemble Clustering Combined with Cluster Optimization (ECCO), an open-source Python UI that provides a fast, scalable framework for ensemble clustering of large-scale data. It includes zero-code integration of novel ensemble clustering methods and many pre- and post-processing functionalities, enabling researchers to efficiently integrate advanced clustering methodologies into their analysis pipelines.

## Introduction

The ongoing omics revolution is dramatically changing biological research. Studies that once featured tens or hundreds of analytes now capture thousands, leading to a breadth of multi-omic analyses of diseases and their underlying mechanisms (***Barapour et al., 2026***; ***Zhou et al., 2024***). Complex diseases such as osteoarthritis are now thought to arise from interactions among multiple biological pathways (***Wilson et al., 2025***). These shifts require researchers to leverage advanced bioinformatic techniques to identify biological mechanisms of disease and quantify disease heterogeneity (***Ronan et al., 2016***). As large-scale multi-omics studies become mainstream, there is a critical need for efficient, validated implementations of the latest analytical methods.

Clustering is an unsupervised machine learning approach that has become a critical part of biological studies (***Oghabian et al., 2014***; ***Chen et al., 2024***; ***Barapour et al., 2026***; ***Hislop et al., 2022***; ***Wallace et al., 2022***; ***Welhaven et al., 2024***). For example, our lab has used clustering to identify novel metabolic endotypes of osteoarthritis (***Wallace et al., 2022***), rapid shifts in synovial fluid metabolism, sex-specific differences in metabolic profiles after joint injuries (***Welhaven et al., 2024***; ***Hislop et al., 2022***), and changes in cortical bone metabolism in chronic kidney disease (***Stauffer et al., 2026***). However, each clustering approach comes with its own set of assumptions and limitations that directly impact the resulting biological insight. This means that an analytical choice could impact a field’s direction rather than the underlying biology. This challenge has been exacerbated by the fact that modern-day omics experiments measure thousands of analytes, leading to the curse of high-dimensionality, where data sparsity and noise mask true biological signals (***Ronan et al., 2016***). To mitigate these challenges, recent studies have suggested using ensemble clustering approaches (***Ronan et al., 2016***). Ensemble clustering works by first determining a set of clustering solutions (2 or more), then aggregating them to determine the analytes that consistently cluster together (***Golalipour et al., 2021***). Before combining the cluster solutions into an ensemble, the number of clusters must be specified or determined using an internal clustering metric (e.g., Average Silhouette Metric (***Rousseeuw, 1987***)). One approach to combining cluster solutions is voting consensus, in which each analyte pair is assigned a weight based on how consistently they cluster together across all cluster solutions. For example, an analyte pair that clusters together in every solution receives a weight of 1 (***Golalipour et al., 2021***). Ensemble clustering has strong potential to broadly improve bioinformatic analysis; however, it has yet to be widely adopted despite the availability of two coding libraries (***Chiu and Talhouk, 2018***; ***Ronan et al., 2018***). A potential explanation for this is the lack of an easy-to-use user interface (UI) and the need for programming expertise to implement. Therefore, we created an easy-to-use open-source Python UI that can be readily utilized by researchers of all backgrounds, with several key improvements:

- Unix and Windows executables allow users to download and run the ECCO UI immediately.
- Codeless generation of custom ensemble cluster solutions.
- User-facing reports on cluster similarity and performance.
- Multi-core processing that scales with available computational resources.
- Various dissimilarity metrics for many types of biological data (Appendix - Figure 1, Appendix - Table 1).
- Built on scipy and scikit-learn, allowing for rapid expansion to other clustering approaches.

ECCO is an open-source Python UI that provides key functionalities for clustering biological data and is scalable, user-friendly, and readily adaptable to researchers’ diverse needs. Users can run ECCO on their local computers or compute clusters, scaling the tool with their available computational resources. In addition, we offer a suite of 24 functionalities, including both pre- and post-processing tools within clustering pipelines and as standalone features (Appendix 1). The current version of the UI (v1.0.3) focuses on agglomerative hierarchical clustering ensemble clustering solutions and is intended to serve as a tool for rapid extraction and analysis of identified analyte clusters. Additionally, this UI serves as a framework for expansion into additional mono-clustering algorithms and ensemble clustering methodologies. Taken together, ECCO provides efficient ensemble clustering analyses and a framework for expansion into a generalizable tool for clustering biological data.

## Methods: ECCO workflow

ECCO is a four-step workflow from data pre-processing to ensemble cluster generation (Figure 1). Users can perform ECCO by submitting an ensemble specification and omics data file (Figure 1, left), and selecting data pre-processing parameters. After submission, ECCO pre-processes the data, generates cluster solutions, and determines the best cluster solution based on the optimal value of an internal cluster metric (e.g., the cluster solution with Average Silhouette Width closest to 1)(Figure 1, middle). Finally, the ensemble cluster solution (i.e., consensus clusters) is calculated (Figure 1, middle), and users are provided with Rand and adjusted Rand index evaluations of each cluster solution, clustergrams of the ensemble cluster solution (Figure 1, right), and output tables of each cluster.

**Figure 1.**
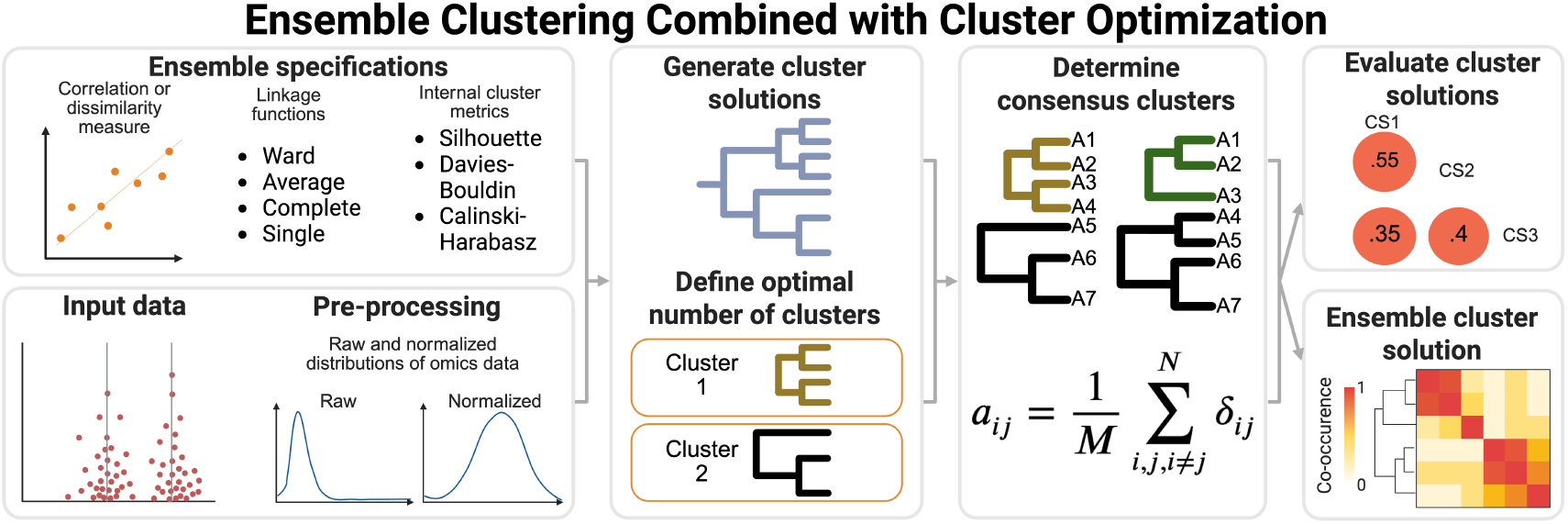
Workflow for Ensemble Clustering Combined with Cluster Optimization. Users input an ensemble specification file and their data, and select pre-processing parameters (left). Cluster solutions are generated and optimized, and the consensus clusters are determined (middle). Cluster evaluation and the resulting ensemble clusters are provided to the user (right).

## Algorithm: Ensemble Clustering Combined with Cluster Optimization

After the input data are pre-processed, the ECCO workflow first generates multiple cluster solutions for the M clustering approaches in the ensemble specification file. Next, internal cluster metrics (Appendix 1) are used to determine the optimal number of clusters for each clustering approach. After which ECCO calculates how often analytes (P) co-occur in clusters across the M cluster solutions, using a voting consensus ensemble approach (***Golalipour et al., 2021***), see equation 1.

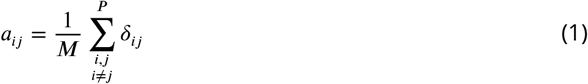

where M is the number of cluster solutions in the ensemble, P is the number of analytes being clustered, and *δ*_*ij*_ is defined as:

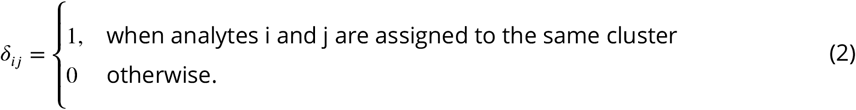

We programmed this ensemble approach such that no adaptation is needed to generate novel clustering ensembles, while also incorporating multi-core processing to improve the efficiency of calculating each cluster solution (the bottleneck of these analyses). Users’ only limitation for ensemble cluster solutions is their available computational resources. Currently, only agglomerative hierarchical clustering of analytes is considered. The ideal number of cluster solutions to consider in the ensemble depends on the input data, and thus, users must select this parameter by considering reasonable expectations of their datasets.

## User Input: Ensemble Specification

The ensemble specification file must contain the input parameters for the dissimilarity metric, linkage function, internal cluster metric, and correlation coefficient or pairwise dissimilarity metric (Table 1) among the P analytes. Users can customize the ensemble by adapting the input parameters in the specification file. The current version of ECCO (v1.0.3) accepts all dissimilarity metrics available within scipy.spatial.distance, and linkage functions of agglomerative hierarchical clustering provided by scikit-learn (***Pedregosa et al., 2011***). Users need to specify an internal cluster metric for each clustering methodology (Table 1) with three currently available: Calinski-Harabasz (CH) (***Caliński and Harabasz, 1974***), Average Silhouette Width (SIL) (***Rousseeuw, 1987***), and Davies-Bouldin Index (DBI) (***Davies and Bouldin, 1979***). To compare observations, users must specify either a Pearson or Spearman correlation coefficient or add pw if they want to leverage a pairwise distance metric (e.g., Euclidean) among the P analytes (Table 1). See Appendix 1 for definitions of the available dissimilarity metrics (Appendix Table 1 and internal cluster metrics.

**Table 1.** Example Specification File for ECCO.

| Dissimilarity | Linkage | Internal Cluster Metric | Correlation |
| --- | --- | --- | --- |
| CNS<sup>a</sup> | Ward | CH<sup>c</sup> | Spearman |
| CS<sup>b</sup> | Ward | CH<sup>c</sup> | Spearman |
| CNS | Complete | SIL<sup>d</sup> | Spearman |
| CS | Complete | SIL<sup>d</sup> | Spearman |
| CNS | Average | DBI<sup>e</sup> | Spearman |
| CS | Average | DBI<sup>e</sup> | Spearman |

<sup>a</sup> CNS: Correlation no square root $1 - |r(Y, Y')|$ <sup>b</sup> CS: Correlation square root $\sqrt{2(1 - |r(Y, Y')|)}$ <sup>c</sup> CH: Calinski-Harabasz Index <sup>d</sup> SIL: Average Silhouette Width <sup>e</sup> DBI: Davies-Bouldin Index

## User Input: Data

Users can input continuous, binary, or count data (Appendix - Table 1), with the only limitation being that the data needs to match our specified structure (Table 2). Specifically, data need to be input as analytes (P) by samples (n) format, with the first column being m.z. (mass-to-charge ratio) and the last column being r.t. (retention time). Both the m.z and r.t. columns contain identifiers and thus can be populated with appropriate labels based on the input data. Any missing values are handled by immediately displaying an error and prompting the user to resubmit the input data without missing values.

**Table 2.** General Format for Input Data Table.

| m.z.<sup>a</sup> | S1<sup>c</sup> | S2 | S3 | S4 | r.t. |
| --- | --- | --- | --- | --- | --- |
|  | G1<sup>d</sup> | G1 | G2 | G2 |  |
| ID1<sup>b</sup> | 1233244.5678 | 23435.6789 | 3436.7890 | 456723.8901 | rt1<sup>e</sup> |
| ID2 | 986.5432 | 876345.4321 | 765344.3210 | 6543095.2109 | rt2 |

<sup>a</sup> m.z. - Mass-to-charge ratio or identifier.
<sup>b</sup> ID#- Analyte identifier.
<sup>c</sup> S# - Sample identifier.
<sup>d</sup> G# - Group identifier.
<sup>e</sup> rt# - Retention time column for mass spectrometry data or additional identifier.

## Example Analysis and Interpretation of ECCO Outputs

In this section, we apply ECCO to a previously analyzed metabolomics dataset (***Carlson et al., 2019***) and provide detailed descriptions of how to interpret the results.

### Case Study: Synovial Fluid Metabolomics Analysis using ECCO

Synovial fluid was collected post-mortem from n=75 patients, with concurrent radiographic evaluation of osteoarthritis (OA, graded by Outerbridge system) (***Carlson et al., 2019***; ***Outerbridge, 1964, 1961***). Metabolites were extracted using an established protocol (***Carlson et al., 2019***; ***Kim et al., 2017***), analyzed using liquid chromatography-mass spectrometry, and the raw data were processed using XCMS (***Domingo-Almenara et al., 2018***). 9962 metabolite features were detected, and 1361 passed median intensity filtering (*i*.*e*. non-zero median intensities in each group) (***Carlson et al., 2019***). Metabolites with zero intensities were those that fell below the mass spectrometer’s limit of detection. Samples were then grouped into no OA, early OA, or late OA groups, but for brevity, here we focus on clustering the metabolites for the 17 samples classified into the late OA group. We set the data pre-processing parameters to log-transformation (base-10), auto-scaling (standardized to z-scale), and specified a viridis colormap. The ECCO UI expects users to handle limit of detection challenges before inputting data, but to ensure robust performance, a negligible positive value is added before log-transforming the analyte intensities. For this example, we input the default ensemble specification file (Table 1), and input the data with 19 columns (m.z, 17 samples, and r.t. columns), and 1361 analytes (rows); no group identifier was specified. On a MacBook Pro with M1 Pro using 6 threads, the ECCO workflow completed in 50 seconds, automatically plotting and saving the following outputs:

- Rand and adjusted Rand index bubble plots comparing each cluster solution (Figure 2A-B) (***Rand, 1971***; ***Hubert, 1985***).
- Rand and adjusted Rand index bubble plots comparing each cluster solution to the ensemble cluster solution (Figure 2C).
- Clustergram of ensemble cluster solution applied to the co-occurrence matrix (Figure 2D).
- Clustergram of the original data with outlines of the clusters identified by ECCO (Figure 2E)
- Spreadsheet containing the co-occurrence matrix.
- Individual spreadsheets for each cluster identified by ECCO, containing the observations present in the cluster.

**Figure 2.**
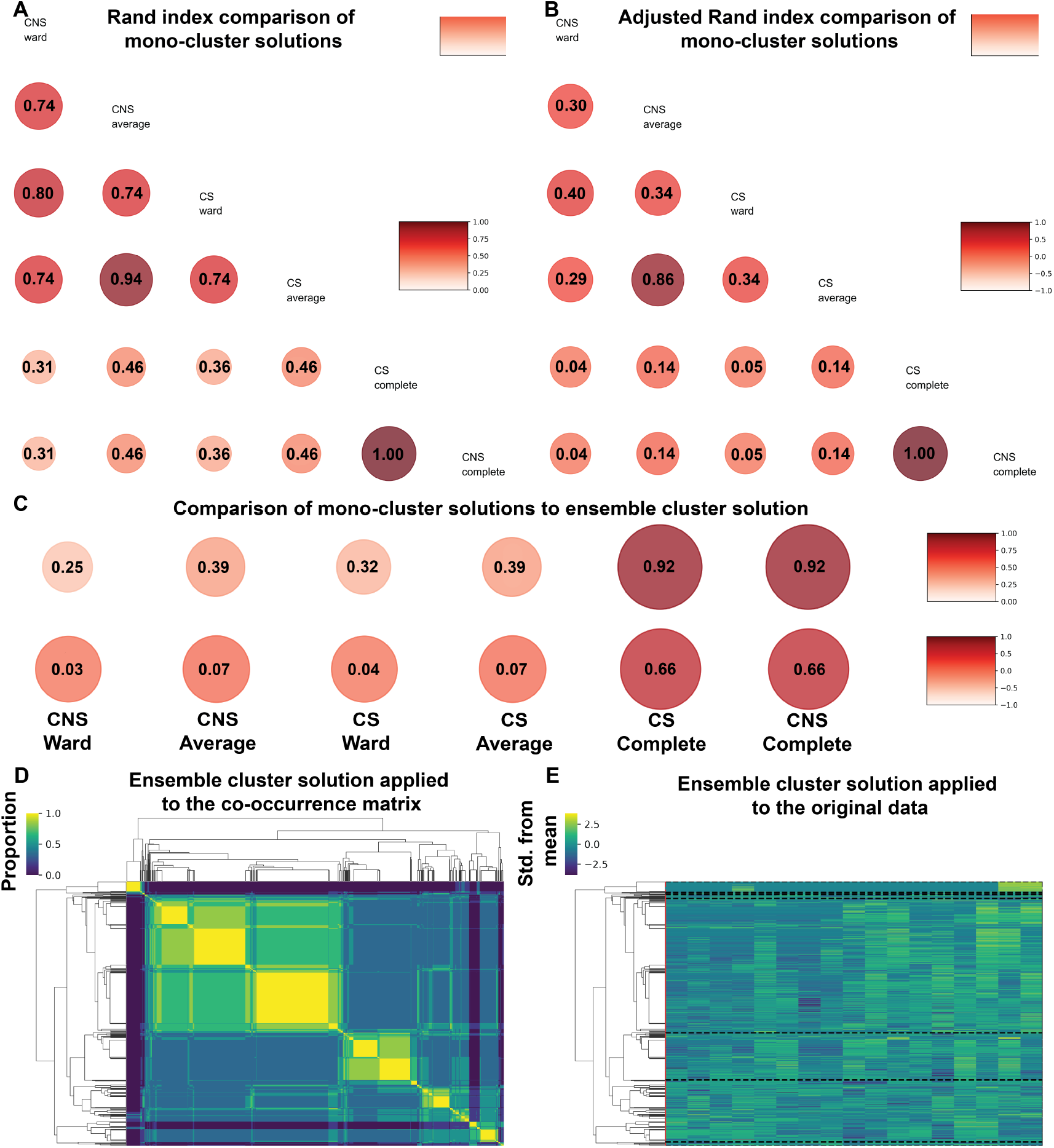
Case Study of Synovial Fluid Metabolism using ECCO. A-B) Comparison of each mono-cluster solution, by Rand-index (A) and adjusted Rand-index (B). C) Comparison of each mono-cluster solution to the ensemble cluster solution using both the Rand-index and adjusted Rand-index. D-E) Clustergrams of the co-occurrence matrix, and the original data showing the clusters identified by ensemble clustering.

### Interpretation of Rand and Adjusted Rand Index

Comparison of mono-cluster solutions (e.g., Complete linkage hierarchical cluster solution) is critical to understanding if similar analytes are consistently clustering. ECCO automatically generates visuals for two well-accepted external cluster evaluation metrics, the Rand index (***Rand, 1971***) and the adjusted Rand index (***Hubert, 1985***). Additionally, external cluster evaluation metrics are also provided in a separate functionality, including the Normalized Mutual Information (***Danon et al., 2005***) and adjusted Normalized Mutual Information (***Vinh et al., 2009***) metrics. The Rand index determines cluster similarity by determining if the cluster solutions agree on the observations that cluster together and those that do not cluster together, with an index of 0 meaning no agreement between cluster solutions, and 1 meaning perfect agreement (***Rand, 1971***). The adjusted Rand index corrects the Rand index for agreement expected by chance (***Hubert, 1985***), with values ranging from −1 to 1, and anything less than 0 representing less agreement on clusters than expected. ECCO represents both the Rand and adjusted Rand index values with increasing bubble size and darkening red color as they go to 1. In our case study, we found that Rand index values varied from 0.31 to 1 (Figure 2A), suggesting cluster solutions ranged from minimal to perfect agreement. For the adjusted Rand index, we found that all mono-cluster solutions achieved greater than random chance agreement, 0.04 to 1 (Figure 2B). These results indicate a high diversity of cluster solutions, which is important to improve ensemble performance (***Golalipour et al., 2021***). We next investigated whether an ensemble cluster solution is necessary by comparing each mono-cluster solution to the ensemble cluster solution (Figure 2C). We found that the complete-linkage clustering solution achieved strong agreement with the ensemble clustering solution, with a Rand index of 0.92 and an adjusted Rand index of 0.66 (Figure 2C). However, when clustering 1362 metabolites, a Rand index of 0.92 would suggest the clusters that >100 metabolites belonged to differed between the clustering solutions, more than enough to impact pathway enrichment results. Therefore, it seems that the ensemble clustering should be used as it combines evidence from multiple clustering solutions.

### Interpretation of Clustergrams

ECCO outputs two clustergrams for visualization of the ensemble cluster solution, one of the cooccurrence matrix and the other of the pre-processed data, with black dashed-line boxes representing the identified clusters (Figure 2D-E). The clustergram (Figure 2D) of the co-occurrence matrix is colored from 0 to 1, with 1 representing metabolites sharing a cluster in each cluster solution. In our case study, we see several distinct yellow (yellow representing 1) squares (Figure 2D), suggesting that, independent of the cluster solution, many metabolites always clustered together. This is consistent with our analysis of the Rand and adjusted Rand index, which suggested that the cluster solutions were identifying similar clusters of observations. Finally, ECCO provides a visualization of differences in metabolic expression between each sample within each of the identified clusters (Figure 2E). In this analysis, we see a cluster at the top of the clustergram where two samples have higher expression of a set of metabolites, whereas nearly all the other samples have minimal expression (Figure 2E, top cluster). ECCO automatically saves each cluster and the co-occurrence matrix from the ensemble solution. For metabolomics analysis, these files can be submitted to the Peaks to pathways functionality (Table 2), which transforms the cluster files for submission to Mummichog (***Li et al., 2013***) for pathway enrichment analysis.

## Implementation Summary and Additional Functionalities

We provide a full list of ECCO functionalities and their inputs and outputs in Table 3 and detailed descriptions of each functionality on our Wiki pages. We expect that users will be able to interact with this tool without needing to write any code. However, the open-source nature of this tool allows users to add additional features, such as novel workflows, that can be readily made into clickable buttons for reusability. For example, we will soon add locally-weighted ensemble clustering (***Huang et al., 2018***) by adding the workflow to the codebase and a button to the UI that calls the workflow.

**Table 3.**
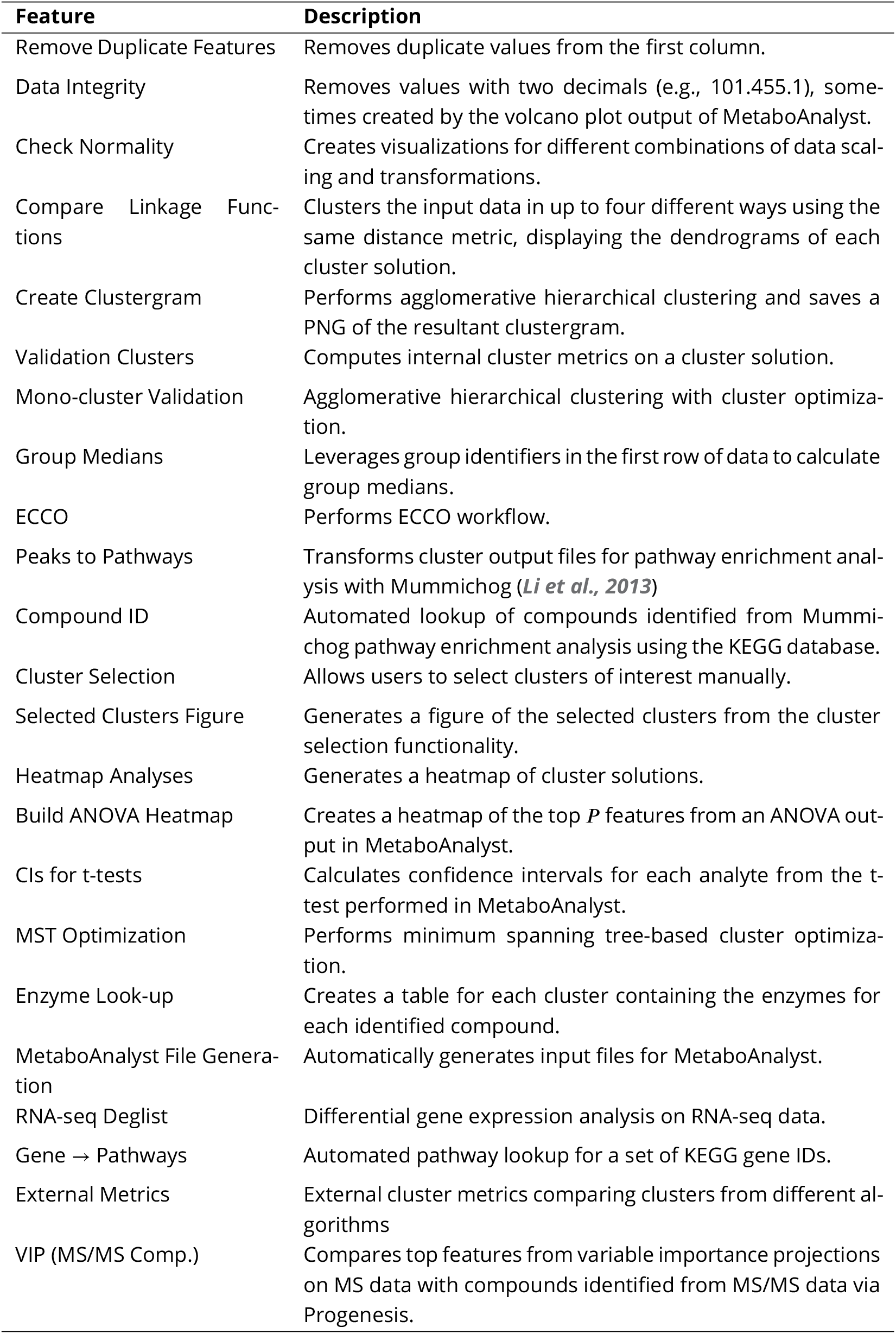
ECCO Functionality Overview.

## Discussion

ECCO is a flexible, multifunctional UI that enables efficient calculation of ensemble hierarchical cluster solutions along with pre- and post-processing capabilities and a suite of additional tools for bioinformatic analyses. The defining functionalities of ECCO are 1) the off-the-shelf usability of the UI, and 2) the ability to generate novel ensembles with zero coding. Users can immediately use ECCO by simply downloading either Unix/Mac or Windows executables from GitHub (v1.0.3). Novel ensembles can be generated by updating a single CSV file, and our open-source codebase allows users to expand the UI’s functionality to incorporate additional non-hierarchical clustering algorithms to meet their needs. Thus, the ECCO UI represents a critical open-source framework for implementing ensemble clustering for biological and non-biological data.

ECCO UI provides a comprehensive suite of tools for data preprocessing, post-processing, clustering, and evaluating internal and external cluster performance, with built-in logging and error handling. The UI handles missing data by automatically displaying an error and prompting the user to resubmit a corrected input file. For customization, each functionality provides users with a series of options. For example, when performing ECCO, users need to select a colormap, data scaling, and data transformation if they want to run a default ensemble (Table 1), and if they want metabolic pathway enrichment files for each cluster that is identified. Users can cluster analytes or samples by simply transposing the input data table from analytes (rows) x samples to samples (rows) x analytes. Detailed explanations of each functionality can be found on our Wiki, and example input and output files are provided for each functionality. All functionalities except ECCO can run locally with reasonable run times; we recommend running ECCO on multiple cores to ensure reasonable compute times.

ECCO has several limitations that will be addressed in future releases. First, our initial release focuses on agglomerative hierarchical clustering and consensus-based ensemble clustering. Future releases will incorporate additional clustering algorithms (e.g., K-means), and other ensemble approaches such as a selective clustering ensemble based on covariance (***Golalipour et al., 2021***). Second, the current workflows are all run through the UI, limiting users to compute clusters with enabled UIs for larger datasets. Given the O((# of analytes)^3^) (***Bataineh, 2022***) complexity of agglomerative hierarchical clustering, to efficiently determine cluster solutions, a high-performance compute cluster is recommended for modern omics applications; thus, incorporating a non-UI workflow would improve usability. Finally, ECCO uses the optimal values of internal cluster metrics (e.g., minimum Davies-Bouldin index) to determine i) the number of clusters for a cluster solution, and 2) the ensemble cluster solution for a predefined number of clusters (100 or fewer). Future versions will allow users to input the number of clusters for each cluster solution, customize the number of clusters to consider, provide visuals of the internal cluster metric results, and incorporate statistical approaches to determining the significance of clusters (e.g., using the Gap Statistic, ***Tibshirani et al. (2001)***). We also plan to expand the UI by incorporating statistical modeling approaches and improving usability. We aim to achieve this by building a community-based coalition to add the latest functionalities to this UI.

## Acknowledgments

This study was funded in part by grants from the NSF (CMMI 2140127) and NIH (NIAMS R01AR073964 and R01AR081489). We would like to thank Hope Welhaven, Erik P. Myers, and Avery Welfley for helping identify bugs in ECCO during development.

## Appendix 1

### Internal Cluster Metrics

The ECCO UI currently supports three internal cluster metrics (***Golalipour et al., 2021***) that quantify the optimal cluster solution. Most internal cluster metrics follow a similar approach, finding the ratios between inter-cluster separation (e.g., separation between cluster centers) and intra-cluster compactness (e.g., the spread of data objects in a cluster). The optimal cluster solution is determined by generating cluster solutions for 2–C clusters (where C ranges from 2 to P-1) and calculating the internal cluster metric for each solution. Depending on the metric, the optimal solution is either a maximum or a minimum. The current metrics include the Average Silhouette Width (***Rousseeuw, 1987***), the Calinski-Harabasz index (***Caliński and Harabasz, 1974***), and the Davies-Bouldin index (***Davies and Bouldin, 1979***).

The Average Silhouette Width (***Rousseeuw, 1987***), determines the optimal number of clusters by:

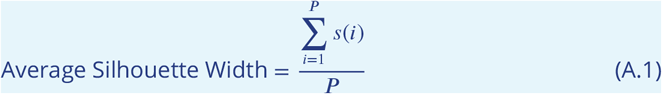

P represents the total number of analytes, and s(i) represents the silhouette widths for each

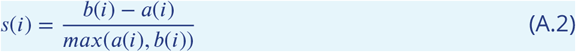

a(i) represents the average dissimilarity between analyte i and analytes in its cluster, and b(i) is the minimum of the average dissimilarity between observation i and the observations in cluster C; s(i) can range from −1 to 1. An Average Silhouette Width of 1 represents an ideal cluster solution.

The Calinski-Harabasz index (***Caliński and Harabasz, 1974***), uses a sum-of-squares-based formulation to determine the optimal cluster solution:

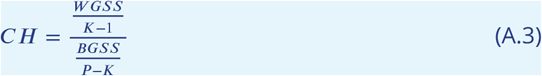

where WGSS is the within-group sum of squares, K is the number of clusters, BGSS is the between-group sum of squares, and P is the number of analytes. The maximum Calinski-Harabasz score represents the optimal cluster solution. The Davies-Bouldin index (***Davies and Bouldin, 1979***) calculates the optimal cluster solution by summing the maximum ratios of intra-cluster spread to cluster separation:

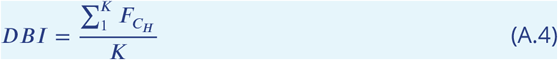

Where K is the number of clusters in the cluster solution 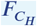 and is:

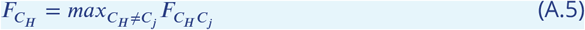

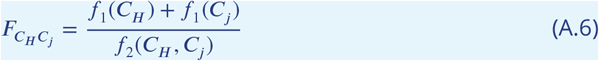

f1 is the average dissimilarity between analytes and their cluster centroids, and f2 is the dissimilarity between centroids. The optimal solution for the Davies-Bouldin index occurs at the minimum value.

### Dissimilarity Measures

The ECCO UI supports all dissimilarity measures available in scipy, and incorporates two correlation-based measures. Typically, correlation-based metrics are transformed to a dissimilarity measure for clustering by either subtracting 1 or using a weighted square root of one minus the correlation (Appendix - Figure 1A-B). However, these transformations create greater dissimilarity to negative correlations, observations that don’t correlate at all, meaning observations with minimal correlation will cluster while those with negative correlations won’t. Thus, we implemented an absolute correlation-based dissimilarity measure (Eq. A.7, 8), which makes positive and negative correlations have similar dissimilarities, while ensuring that analytes with zero correlation do not cluster (Appendix - Figure 1.

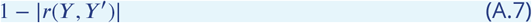

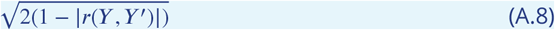

**Appendix 1—figure 1.**
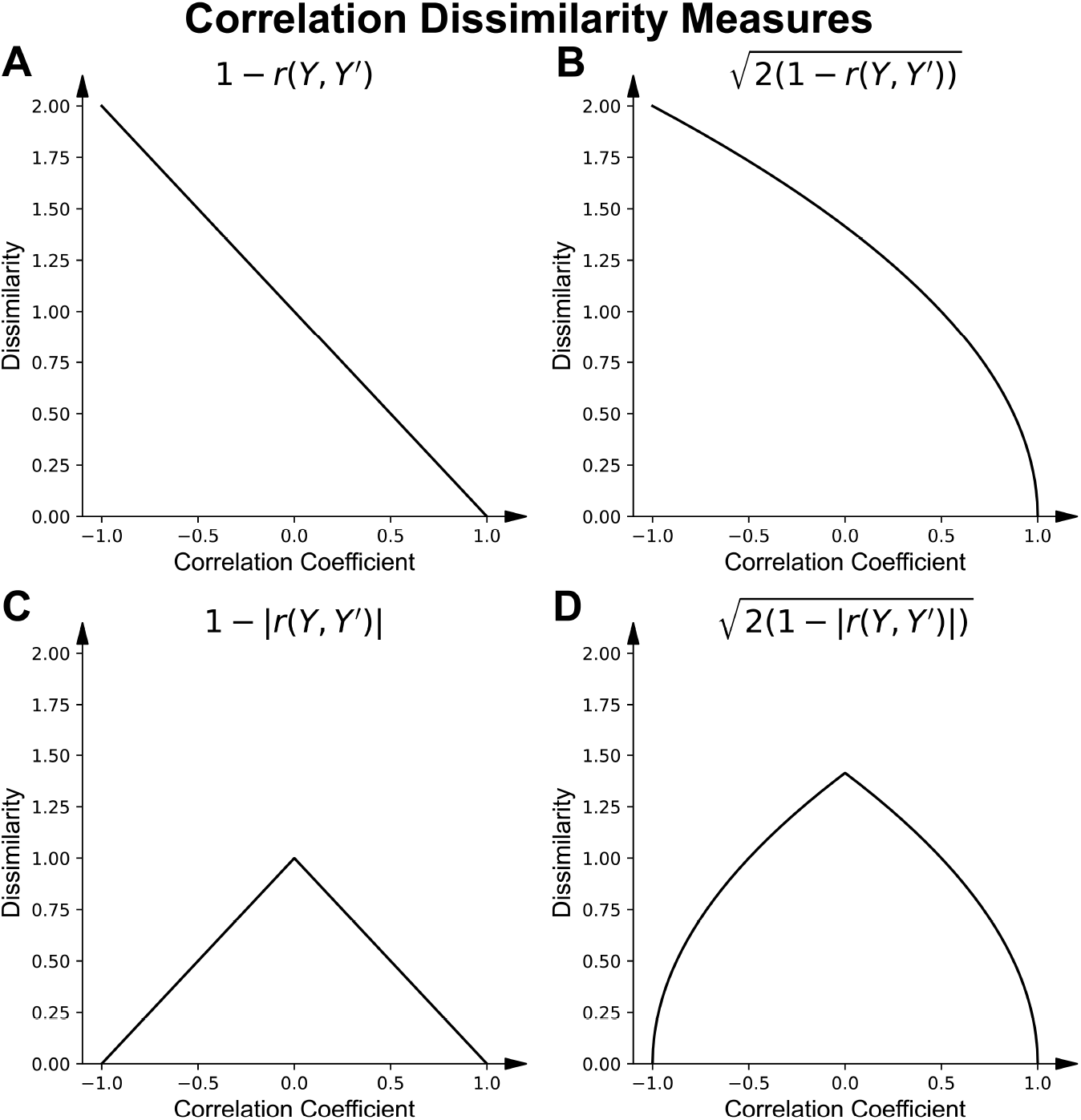
Comparison of correlation dissimilarity metrics. A) Correlation dissimilarity B) Weighted square-root correlation dissimilarity. C) Absolute Correlation dissimilarity. D) Weighted square-root absolute correlation dissimilarity.

**Appendix 1—table 1.** Dissimilarity Metrics and Data Types.

| Dissimilarity | Data Type | Example Data |
| --- | --- | --- |
| Manhattan | Continuous | Mass Spectrometry Intensity data |
| Euclidean | Continuous | Mass Spectrometry Intensity data |
| Chebyshev | Continuous | Mass Spectrometry Intensity data |
| Cosine | Continuous | Mass Spectrometry Intensity data |
| Bray-Curtis | Counts | Shotgun Metagenomics, RNA-seq |
| Jaccard | Binary | Shotgun Metagenomics, RNA-seq |
| Hamming | Sequences | Genetic sequences |
| Jensen-Shannon | Probability Distributions | Microbiome data |
| Canberra | Continuous | Mass Spectrometry Intensity data |
| Yule | Binary | Shotgun Metagenomics, RNA-seq |

